# Independence between pre-mRNA splicing and *de novo* DNA methylation in an isogenic minigene resource

**DOI:** 10.1101/124073

**Authors:** Kyster K. Nanan, Cody Ocheltree, David Sturgill, Mariana D. Mandler, Maria Prigge, Garima Varma, Shalini Oberdoerffer

## Abstract

Actively transcribed genes adopt a unique chromatin environment with characteristic patterns of enrichment. Within gene bodies, H3K36me3 and cytosine DNA methylation are elevated at exons of spliced genes and have been implicated in the regulation of pre-mRNA splicing. H3K36me3 is further responsive to splicing, wherein splicing inhibition led to a redistribution and general reduction over gene bodies. In contrast, little is known of the mechanisms supporting elevated DNA methylation at actively spliced genic locations. Recent evidence associating the *de novo* DNA methyltransferase Dnmt3b with H3K36me3-rich chromatin raises the possibility that genic DNA methylation is influenced by splicing-associated H3K36me3. Here, we report the generation of an isogenic resource to test the direct impact of splicing on chromatin. A panel of minigenes of varying splicing potential were integrated into a single FRT site for inducible expression. Profiling of H3K36me3 confirmed the established relationship to splicing, wherein levels were directly correlated with splicing efficiency. In contrast, DNA methylation was equivalently detected across the minigene panel, irrespective of splicing and H3K36me3 status. In addition to revealing a degree of independence between genic H3K36me3 and DNA methylation, these findings highlight the generated minigene panel as a flexible platform for the query of splicing-dependent chromatin modifications.

## INTRODUCTION

Genes in higher eukaryotes are characterized by short exons separated by relatively long introns (1,2). To facilitate ligation of contiguous exons, spliceosome assembly occurs co-transcriptionally, while the nascent transcript remains tethered to the chromatin template [(3-6) and reviewed in (7)]. Cotranscriptional splicing further allows for coupling between the spliceosome and intragenic chromatin. Genome-wide studies provide support for such an association, wherein specific chromatin modifications are enriched at exons relative to introns of expressed genes (8,9). In particular, trimethylation of histone H3 lysine 36 (H3K36me3) and methylation of cytosine at the fifth carbon (5- methylcytosine, 5mC) are elevated at spliced exons, and reciprocally depleted at intronless genes and pseudoexons (10-12). These associations persist after correcting for the overall increase in nucleosome occupancy and CpG frequency at exons. Combined with evidence that alternative exons that are included in spliced mRNA are characterized by increased H3K36me3 and 5mC as compared to their excluded counterparts, the distribution of chromatin modifications within gene bodies suggests a functional relationship to splicing (13-15). Indeed, studies of alternative splicing implicate H3K36me3-associated chromatin binding proteins in pre-mRNA splicing decisions through the direct recruitment of splicing factors (16,17). In addition, we and others have shown that methylation of DNA modulates splicing decisions through differential binding of methyl-sensitive transcription factors that obstruct pol II elongation and thereby favor spliceosome assembly at available weak splice sites (14,18,19). These demonstrations raise the possibility that exonic chromatin modifications aid the spliceosome in discerning between productive exon-flanking splice sites and a kaleidoscope of undesired cryptic sites. However, the mechanisms supporting selective enrichment of chromatin modifications at exons remain unclear.

Recent studies of H3K36me3 begin to reveal a reciprocal role for the spliceosome in the establishment of genic chromatin. A hallmark of transcription, trimethylation of H3K36 is catalyzed by the SETD2 methyltransferase via interaction with hyperphosphorylated pol II (20-23). Whereas association with pol II would imply uniform distribution over gene bodies, enrichment of H3K36me3 at exons suggested an active mechanism for discriminating sequences corresponding to exons from the surrounding landscape (24). As a key interpreter of splicing signals, the spliceosome itself emerged as a likely candidate. Indeed, inhibition of splicing through chemical or mutational means resulted in a general reduction in genic H3K36me3 and an overall repositioning to the 3’ ends of genes (10,25). However, it should be noted that splicing is not required for SETD2 function: H3K36me3 is abundantly detected at regions of active pol II transcription in organisms that possess few, if any, introns (26). Together, these studies confirm that splicing “augments” H3K36me3 through a currently unknown mechanism.

Whereas genic DNA methylation displays a similar pattern of enrichment at spliced exons to H3K36me3, the overall dependency on splicing has not been explored. In mammalian genomes, 5mC is primarily found in a symmetric context on CpG dinucleotides, with greater than 70% of such sites modified in human cells (27). Once established by the *de novo* DNA methyltransferases (DNMT3A/B in mammals), methylation patterns are preserved by maintenance methyltransferases (DNMT1 in mammals) that recognize hemimethylated DNA and ensure site-specific propagation in the newly synthesized strand (28). In contrast to promoter methylation, intragenic DNA methylation is positively associated with gene expression (29-31). Like H3K36me3, this association extends to the per exon level, wherein exons that are included in spliced mRNA show elevated 5mC (11,12). Exonic enrichment of DNA methylation is conserved across eukaryotic evolution and is often most strikingly observed in organisms with an overall lower level of methylation (32,33). For example, the <1% of CpGs that are methylated in the honeybee *Apis mellifera* occur almost exclusively within exons of spliced genes (15). When considering the human genome, whole-genome bisulfite sequencing (WGBS) reveals abrupt transitions in methylation at exon-intron junctions, wherein a sharp spike is detected at 5’ splice sites and sharp dip at 3’ splice sites (34) suggesting mechanisms of active targeting.

Given the overall similarity in distribution of H3K36me3 and 5mC within gene bodies, it is tempting to envision a common mechanistic basis (11,12,15,35). The presence of an H3K36me3-interacting PWWP domain in the *de novo* methyltransferases DNMT3A and DNMT3B provides such a link (36,37). Accordingly, ablation of either SETD2 or the PWWP domain resulted in decreased Dnmt3B interaction with genic DNA and an overall reduction in gene body DNA methylation (38,39). While revealing H3K36me3 as a potential scaffold for subsequent DNA methylation, 5mC is notably distinguished from H3K36me3 on several fronts. For example, whereas H3K36me3 is omnipresent across eukaryotic evolution, 5mC is entirely lacking in select species including several prominent model organisms (40-42). Importantly, reintroduction of Dnmt3 enzymes into null contexts results in overall low levels of *de novo* methylation, suggesting that H3K36me3 is an inefficient scaffold at best (38,39). Finally, unlike H3K36me3, which is restricted to expressed gene bodies, 5mC is ubiquitous at intergenic regions, including transcriptionally repressed promoters and intergenic repetitive elements (27,33,43). Together, these results suggest a substantive but nuanced relationship between genic H3K36me3 and 5mC.

Here we report the generation of an isogenic resource for assessing the direct role of splicing in the regulation and impact of intragenic DNA methylation. A panel of minigenes modeled on naturally occurring single-isoform spliced and unspliced contexts were integrated into a single FRT site in 293 T-REx cells. Prior to integration, the minigenes were modified to produce a range of splicing efficiencies. The presence of an inducible promoter upstream of the minigene DNA further allowed for uncoupling of transcription and splicing-dependent effects on the adopted chromatin structure. Examination of the minigene chromatin recapitulated the previous association to H3K36me3, wherein the highest levels were detected in the presence of the most efficiently spliced minigene. Likewise, as would be expected given the low *de novo* methyltransferase expression in 293 cells, stable introduction of the unmethylated minigenes resulted in basal hypomethylation. However, exogenous expression of the DNMT3 enzymes yielded equivalent gains in DNA methylation, irrespective of transcriptional activity, splicing potential, or H3K36me3 status. These findings reveal a degree of independence in the mechanisms guiding emerging H3K36me3 and 5mC at transcriptionally active gene bodies and establish the generated isogenic panel as a valuable resource for elucidating the respective contributions of transcription and splicing on genic chromatin.

## MATERIAL AND METHODS

### Identification of model single-exon and two-exon minigenes

Annotation tables for mm9 (mouse) were retrieved via the UCSC Table Browser (https://genome.ucsc.edu/cgi-bin/hgTables). Genes were filtered by exon count and mammalian conservation to generate a list of single- or two-exon genes that were largely sequence-conserved between mouse and human. Next, genes with more than one isoform and a length greater than 1 kb from transcriptional start site (TSS) to transcriptional termination site (TTS) were removed to generate a list of potential minigene candidates. The remaining genes were screened using the online Splice Site Analyzer Tool (http://host-ibis2.tau.ac.il/ssat/SpliceSiteFrame.htm) to identify candidate minigenes with the fewest and weakest cryptic splice sites.

### Molecular cloning of plasmid constructs

All minigene DNA sequences were commercially synthesized by GenScript (Piscataway, NJ, USA) and inserted into the pUC57-simple vector by the manufacturer. Each minigene sequence was flanked by 5’ KpnI and 3’ BamHI restriction sites, with the exception of the *Tnp1* cDNA minigene, which was flanked by 5’ and 3’ KpnI sites. Minigenes were inserted into the KpnI/BamHI sites (KpnI site for *Tnp1* cDNA) of pcDNA5 FRT/TO (Invitrogen) using standard molecular cloning techniques. The *Tnp1* gDNA was modified by deleting the intron or mutating consensus and cryptic splice sites to generate the *Tnp1* cDNA or *Tnp1* ΔSS minigenes, respectively. *Tnp1* ΔSS+5’3’SS was generated by restoring the consensus splice sites of the *Tnp1* ΔSS minigene to their wildtype sequences. The *Pfn3* ORF was modified to generate *Pfn3*+5’3’SS by the addition of a consensus splice acceptor at its 5’ end (immediately upstream of the start codon) and splice donor towards the 3’ end. Complete minigene sequences are provided in FASTA format in the online Supplementary Materials (Figure S1B).

### Cell culture

Parental Flp-In 293 T-REx cells (Invitrogen) were cultured in complete DMEM (Gibco) containing 10% tetracycline-reduced FBS (Clontech), 2 mM L-glutamine (Gibco) supplemented with 5 μg/ml blasticidin, and 400 μg/ml Zeocin (Invivogen). Minigene integration was achieved according to the manufacturer’s instructions. Briefly, Flp-In 293 T-REx cells were transfected (Lipofectamine 2000, Invitrogen) with minigene-encoding pcDNA5 FRT/TO and Flp recombinase-encoding pOG44 (Invitrogen). 48-hours post-transfection, targeted cells were selected through addition of 5 μg/ml blasticidin and 100 μg/ml Hygromycin B Gold (Invivogen). Genomic DNA was extracted and PCR was performed with a minigene-derived forward primer and insertion-site-derived reverse primer to confirm site-specific integration. Gene expression was induced with 1 μg/ml doxycycline (Sigma) for up to 24 hours, as indicated. To achieve DNMT3 overexpression, host cells were transfected with equimolar pCMV-Sport6-hDNMT3A (Thermo Scientific, #6150112) and pCMV-Sport6-mDnmt3l (Thermo Scientific, #30447906) or pcDNA3.1-myc-hDNMT3B (Addgene, #35522) with Lipofectamine 2000, according to the manufacturer’s instructions.

### Western blotting

Immunoblotting for protein expression was performed with standard techniques. Briefly, cells were lysed in NP-40 lysis buffer (1% NP40, 150 mM NaCl, 50 mM Tris-HCl; pH 8.0) and 50 μg protein lysates (Bio-Rad, #5000201) were resolved by SDS-PAGE gel and transferred to PVDF (Millipore, #IPVH00010). The following antibodies were used for immunoblotting: DNMT3A (Cell Signaling Technology, #2160S), DNMT3B (Cell Signaling Technology, #D7070), DNMT3L (abcam, #ab3493), β-tubulin (Cell Signaling Technology, #2146S), and HRP-conjugated anti-rabbit antibody (Cell Signaling Technology, #7074). Membranes were exposed to X-ray film after brief incubation with chemiluminescence reagents (GE, #RPN2106).

### Southern blotting

Genomic DNA was extracted from minigene cell lines, digested to completion with EcoRV, resolved on a 0.8% agarose gel, and transferred to a nylon membrane. DNA was crosslinked to the membrane by UV irradiation (Stratagene) and blocked with prehybridization buffer (6X SSPE, 100X Denhardt’s Solution, 1% SDS, 50 μg/ml sheared, single-stranded salmon sperm DNA) for 2 hours at 65 °C. The blocked membrane was then incubated with radiolabeled probe against the *lacZeo* cassette (Prime It II Random Primer Labeling, Agilent Technologies) in hybridization buffer (6X SSPE, 1% SDS, 100 μg/ml sheared, single-stranded salmon sperm DNA) at 65 °C with gentle rotation. After overnight incubation with probe, the blot was washed (1X SSPE, 0.2% SDS) and exposed to a phosphor screen.

### RNA isolation and analysis

For quantitative- and radioactive-RT-PCR, RNA was purified from 293 T-REx cells using RNeasy mini columns (QIAGEN); DNase treatment was performed on-column and post-elution using RNase-Free DNase set (QIAGEN) and TURBO DNA-free DNase (Ambion) reagents, respectively. 500 ng of RNA was utilized for cDNA conversion with Superscript III reverse transcriptase (Invitrogen) and primed with random hexamers, except where indicated, according to the manufacturer’s instructions. For LightCycler (Roche) quantification of minigene expression, cDNA was amplified in the presence of SYBR Green (Roche) with primers against a specific exon and normalized to a panel of 3 endogenous reference genes (*ALDOLASE, GAPDH*, and *RPS16*) using the ΔCT method: relative expression = 2^(mean CT_*ref*_. − CT_*minigene*_). For splicing analysis, template cDNA was amplified by *Taq* polymerase (NEB) using *Tnp1*-specific intron-flanking primers, along with control primers against the reference gene *TUBB*, in a multiplexed PCR reaction containing 33 nM α-^32^P-dCTP (3000 Ci/mmol, Perkin Elmer). PCR fragments were separated by electrophoresis on a 5% TBE-polyacrylamide gel, which was then dried, and exposed to a phosphor screen. For the single-exon *Pfn3* minigene, radioactive RT-PCR was performed using forward and reverse primers directed against the 5’ and 3’ ends of the gene, respectively.

RNA used in 3’-end RT-PCR was extracted using Trizol and poly(A) site usage was determined as described in Elkon *et al*. (44). Briefly, 1 μg of total RNA was dissolved in fragmentation buffer (100 mM Tris, pH 8.0, 2 mM MgCl_2_), heated to 95 °C for 3 minutes, then immediately chilled on ice. Conversion to cDNA was achieved with M-MLV (Sigma) and a hybrid P7/anchored oligo(dT)_25_ primer (5′-AAGCAGAAGACGGCATACGAGATTTTTTTTTTTTTTTTTTTTTTTTTVN-3′), followed by PCR amplification (*Taq* polymerase, NEB) with a gene-specific forward primer and P7 reverse primer. 3’- end RT-PCR products were separated and visualized using standard agarose gel electrophoresis techniques. Primers for gene expression analysis are shown in Table S1.

### Chromatin immunoprecipitation

Crosslinking was performed at room temp with 1% formaldehyde (Sigma) for 5 minutes and quenched with 125 mM glycine (ICN Biomedical) for 10 minutes. Cell membranes were lysed using cold NP-40 buffer (1% NP40, 150 mM NaCl, 50 mM Tris-HCl; pH 8.0) and nuclei collected by centrifugation at 12,000xg for 1 minute at 4 °C. Nuclear pellets were resuspended in ChIP sonication buffer (1% SDS, 10 mM EDTA, 50 mM Tris-HCl; pH 8.0), supplemented with Halt protease inhibitors (Thermo Scientific), and chromatin sheared to an average size between 150 and 500 bp by sonication (Bioruptor Twin, Diagenode). Chromatin preparations were cleared by centrifugation at 20,000xg for 10 minutes at 4 °C and protein content measured by BCA assay (Thermo Scientific). 250 μg of chromatin was diluted at least 10-fold in ChIP dilution buffer (1.1% Triton X-100, 0.01% SDS, 167 mM NaCl, 1.2 mM EDTA, 16.7 mM Tris-HCl; pH 8.1), the appropriate antibody was added, and samples were incubated overnight at 4 °C with rotation. Immune complexes were captured using protein G (Millipore, #16-201) or protein A/G (SantaCruz Biotechnology, #sc-2003) agarose beads blocked with BSA and salmon sperm DNA, and washed with ChIP dilution buffer. Washed beads were resuspended in crosslink reversal buffer (50 mM EDTA, 10 mM Tris-HCl; pH 8.0) and incubated at 95 °C for 10 minutes. DNA was purified by column purification (QIAGEN) and qPCR was performed using SYBR Green chemistry (Quanta or Roche) and gene-specific primers. H3K36me3 enrichment was determined relative to pan-histone H3 [2^(CT_pan-H3_-CT_H3K36me3_)] and normalized to a control amplicon in the vector-introduced and non-expressed Amp gene to correct for variations in IP efficiency. Pan-histone H3 occupancy was determined relative to input [% input = 100*2^^^(CT_input_-CT_IP_)] and normalized to H3 levels at a control locus in the Amp gene. Each ChIP-qPCR experiment was performed on three biological replicates and the data were plotted as mean ± SEM, unless otherwise noted. The following antibodies were used for ChIP: histone H3 (Cell Signalling Technology, #2650; 4 μl per IP), H3K36me3 (abcam, #ab9050; 2 μg per IP), normal rabbit IgG (Cell Signaling Technology, #2727; 2 μl per IP). Primers used for ChIP-qPCR analysis are presented in Table S1.

### Methylation-sensitive restriction enzyme-PCR (MSRE-PCR)

Genomic DNA was purified from minigene host 293 T-REx cells using phenol:chloroform (Ambion, #17907) and ethanol precipitation. For MSRE-PCR, 100 ng of genomic DNA was digested overnight with the methyl-sensitive restriction enzyme HpaII (NEB, #R0171S) or methyl-insensitive enzyme MspI (NEB, #R0106S). Digested DNA was amplified by PCR using *Taq* polymerase (NEB) and primers that flanked HpaII/MspI sites (“methyl-sensitive”) or control sites (“methyl-insensitive”) in *Tnp1* and *Pfn3* minigenes and products were visualized by agarose gel electrophoresis. For quantitative evaluation of DNA methylation by MSRE-qPCR, DNA was digested as described above and amplified using methyl-sensitive and methyl-insensitive primers with SYBR green-based PCR reagents (Roche). Relative DNA methylation levels were calculated according to: 100*2^(Ct_methyl-insensitive_ − Ct_methyl-sensitive_). Where indicated, minigene expression was induced via 24-hour treatment with doxycycline (1 μg/ml final concentration). Data are plotted as mean ± SD for two independent, biological replicates. MSRE-PCR and MSRE-qPCR primer sequences are presented in Table S1.

### Bisulfite-pyrosequencing

Genomic DNA was extracted and purified using either phenol:chloroform extraction (Ambion, #17907) and ethanol precipitation or via the Invitrogen PureLink Genomic DNA Mini kit (Life Technologies), according to the manufacturer’s instructions. Bisulfite conversion and pyrosequencing were performed at the NCI Advanced Technology Research Facility (ATRF). Briefly, 500 ng genomic DNA was bisulfite-treated and purified using a commercial kit (Qiagen, #59104). PyroMark Assay Design Software 2.0 was used to design the assay and pyrosequencing was performed on the PyroMark Q96 MD. Data are presented as a heatmap of individual CpG methylation percentages or summarized as average methylation across the sequenced region; error bars indicate SEM. Primers used for bisulfite-pyrosequencing are listed in Table S1.

### Bioinformatics analysis

#### Gene set definitions

Gene definitions were retrieved from the UCSC canonical genes table, which presents a canonical isoform for each locus (usually the longest), from the UCSC genome browser (45). The canonical isoform was used to define single/multi exon genes. A list of histone genes was retrieved from the HUGO Gene Nomenclature Committee, to filter histone genes from the analysis (46). Genes annotated on chrM or chrY were also removed from analysis.

#### RNA-Seq analysis

For K562, RNA-seq reads were obtained from ENCODE (Experiment number ENCSR000EYO, GEO accession GSM958729). One replicate of this experiment was used (fastq accessions ENCFF000DWV and ENCFF000DXN). Polyadenylated RNA from K562 cell was sequenced, generating ∼70 million paired-end 76bp reads (35 million pairs). T-cell RNA-seq data used in this study were previously described (GSM1936639) (19). Reads were aligned with Tophat v.2.0.10 using Bowtie v1.0.0, reporting one alignment per read (the -g 1 option) (47,48). Gene level expression was quantified (in units of FPKM - expected fragments per kilobase per million reads) using Cufflinks v2.1.1 against the UCSC canonical gene set (47). Genes were partitioned into “expressed” and “non-expressed” with a cutoff of FPKM=1 from these results. For K562, this procedure was run in parallel on additional replicates of RNA-seq data, including a separate library construction, and gene level expression measurements were compared to ensure that this sample was representative (data not shown). All *inter-se* comparisons for K562 RNA-seq yielded a Pearson’s r >= 0.99.

#### ChIP-Seq and MedIP-Seq analysis

ChIP-seq data were batch downloaded from the ENCODE data portal (http://www.encodeproject.org). ChIP data were from K562 cells. Data were downloaded in bigwig format. ENCODE file accessions for each sample used are: H3K36me3 (ENCFF001FWL), H3K4me3 (ENCFF000VDQ), H3K9ac (ENCFF000BYN), H3K27ac (ENCFF000BWY), H3K79me2 (ENCFF000BYH), H3K27me3 (ENCFF000VDE), TBP (ENCFF000ZBQ), and Pol II (ENCFF000YWS). MedIP-Seq experiments were performed as previously described (GSM1936634) (19). Sequencing libraries were constructed from DNA samples with the Illumina TruSeq ChIP Sample Prep Kit (IP-202-1012-1024). Libraries were sequenced on an Illumina NextSeq instrument for 76 cycles in single-end mode. Reads were mapped to the human genome with Bowtie2 v.2.1.0 with default parameters, producing output in SAM format. The default parameters allowed at most one alignment per read (48). The reference assembly used was hg19 from UCSC, via iGenomes (Illumina, San Diego CA). Coverage over gene features was calculated using deepTools, over reference points (TSS) or scaled regions (gene bodies) (49). Heatmaps were generated with the annotated heatmap function in the Bioconductor non-negative matrix factorization (NMF) package (50).

## RESULTS

### Isogenic cell lines for inducible “natural” minigene expression

To examine the impact of splicing on intragenic chromatin, we generated minigene constructs of varying splicing potential. Classical minigenes typically encompass 2 to 3 “test” exons with intervening intronic sequences. As introns are relatively extended as compared to exons, with respective average lengths of approximately 5 kb and 150 bp in the human genome (51), introns are often truncated to only include the ∼200 nucleotides flanking the 5’ and 3’ splice sites (52). Consequently, distal regulatory sequences may be omitted from minigene constructs, resulting in reduced splicing efficiency or the utilization of cryptic splice sites. To generate conditions for clear assignment of spliced versus unspliced states, we mined the genome to identify candidates that naturally meet optimal minigene guidelines. Specifically, we searched for conserved genes that produce a single isoform in both human and mouse cells, that are less than 1 kilobase in length from transcription start (TSS) to termination site (TTS), and that possess a minimal number of potential cryptic sites that could be activated after canonical splice site mutation (Fig. 1A, Table S2). Candidate two-exon, *Tnp1*, and single-exon genes, *Pfn3*, of comparable length were selected to represent splicing competent and deficient contexts, respectively (Fig. 1B).

**Figure 1.**
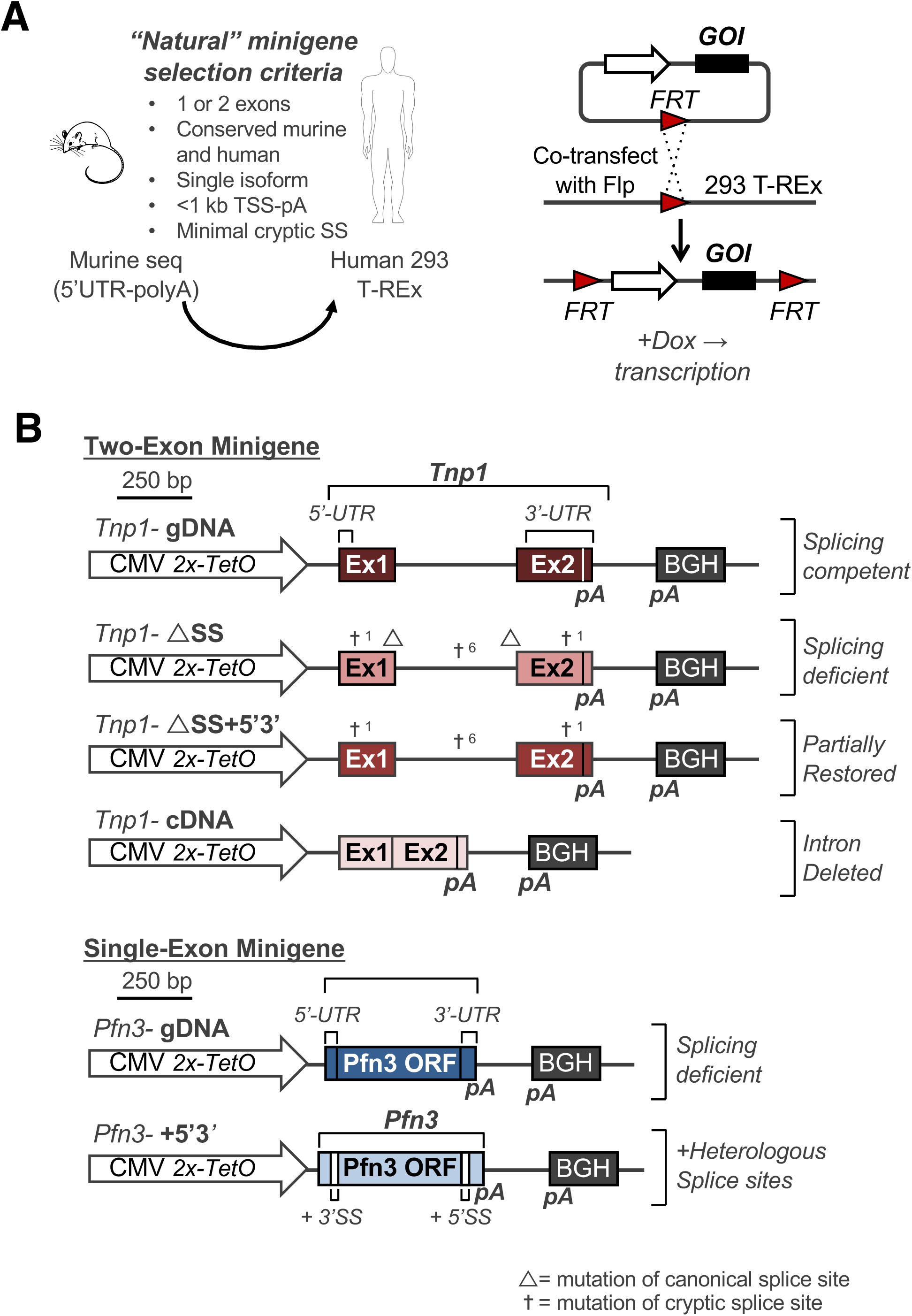
Selection of “natural” single- and two-exon minigenes. **(A)** Murine single-exon and two-exon genes were selected based on several optimal criteria for minigene generation **(left)**. The corresponding sequences were integrated into a single FRT site in human Flp-In 293 T-REx cells via Flp-mediated recombination. Minigene expression is induced through addition of the tetracycline analog, doxycycline (Dox) (right). **(B)** Schematic representations of utilized two-exon (*Tnp1*) and single-exon (*Pfn3*) minigenes. The wildtype *Tnp1* gDNA minigene was modified to generate splicing deficient contexts through mutation of canonical and cryptic splice sites (Tnp1-ΔSS) or intron deletion (Tnp1-cDNA). Re-introduction of only the canonical splice sites to the ΔSS sequence is denoted as *Tnp1*-ΔSS+5’3’. The wildtype *Pfn3* gDNA minigene was modified through addition of heterologous splice sites to create the *Pfn3*+5’3’ minigene. TSS, transcriptional start site; pA, polyadenylation site; SS, splice site; Flp, flippase; FRT, flippase recognition target; ORF, open reading frame.

While minigene analysis of splicing is typically performed in transient transfection, to assess the impact of splicing on genic chromatin, we pursued stable integration into chromosomal DNA. To ensure that all minigenes are exposed to a uniform surrounding chromatin landscape, we utilized <u>F</u>lp <u>R</u>ecombination <u>T</u>arget (FRT)-mediated integration of the minigene DNA into a single genomic location. Briefly, the entire murine genic sequences, including untranslated regions (UTRs) and polyadenylation (p(A)) sites, were cloned into a CMV driven expression vector containing a single FRT site. The splicing competent *Tnp1* minigene (gDNA) further served as template for the generation of splicing deficient contexts through deletion of the intervening intron (cDNA) or mutation of the canonical and predicted cryptic splice site (ΔSS) (Fig. 1B) (53). As variations in DNA GC content can influence chromatin assembly (54-56), conservative dinucleotide substitutions were employed to achieve splice site ablation (i.e. GT to CA, or AG to TC). In addition, the ΔSS minigene was partially “rescued” through restoring the canonical intron-flanking splice sites, while leaving the cryptic substitutions in place (ΔSS+5’3’) (Fig. 1B). Finally, the single exon *Pfn3* minigene was further adapted through the introduction of consensus 3’ and 5’ splice sites upstream and downstream of the *Pfn3* open reading frame (ORF), respectively. The individual minigenes were co-transfected along with plasmid encoding Flp recombinase to drive insertion into a single FRT site in human 293 T-REx cells, as validated through Southern blot (Fig. 1A, Fig. S1).

To confirm that the integrated minigenes were expressed and spliced as expected, total RNA was harvested pre- and post-induction of expression through 24 hours of doxycycline (Dox). 1 μg/ml Dox was selected as an optimal concentration for minigene induction (Fig. S2A). Comparison of reverse transcription performed with oligo(dT) versus random hexamers revealed preferential detection of the spliced mRNA in the oligo(dT) primed cDNA (Fig. S2B), as would be expected based on the known functional link between splicing and polyadenylation (57-59). Thus, to ensure against biased detection of spliced transcripts in the minigene characterization, reverse transcription (RT) was performed with random hexamers. Quantitative assessment of minigene induction in response to 4 hours of Dox was achieved through qRT-PCR with primers against the 3’ end of each transcript (Fig. 2A). Consistent with the reported impact of splicing on transcription (60-64) the *Tnp1* gDNA minigene showed the highest fold induction. Nevertheless, all minigenes were significantly induced and, with the exception of *Tnp1* gDNA, achieved comparable expression, (Fig. 2B). To additionally establish minigene splicing patterns, PCR amplification was performed on cDNA generated following 4 hours of induction in the presence of radiolabelled dCTP α^32^P with *Tnp1* exon 1 forward and exon 2 reverse primers, or with primers against the 5’ and 3’ ends of the *Pfn3* coding sequence (Fig. 2A). Phosphorimaging analysis revealed the desired *Tnp1* splicing outcomes: the gDNA construct was efficiently spliced postinduction whereas the splicing-deficient ΔSS and cDNA constructs produced a single, unspliced isoform (Fig. 2C). As might be expected based on the remaining “cryptic” site substitutions that could potentially impact splicing enhancers, reconstitution of the canonical splice sites in the ΔSS+5’3’SS construct did not fully restore wildtype splicing levels, thus allowing for examination of a partially spliced state in downstream analyses (Fig. 2C). With respect to the *Pfn3* minigenes, both the gDNA and +5’3’SS constructs showed no evidence of splicing (Fig. 2C). The latter is expected given the reversed orientation of 3’ and 5’ splice sites as upstream and downstream of the ORF, respectively. In sum, examination of the minigene-derived transcripts revealed the expected induction and splicing patterns, thereby highlighting comparison of single-versus multi-exonic genes as a valid approach for assessing the impact of splicing on intragenic chromatin.

**Figure 2.**
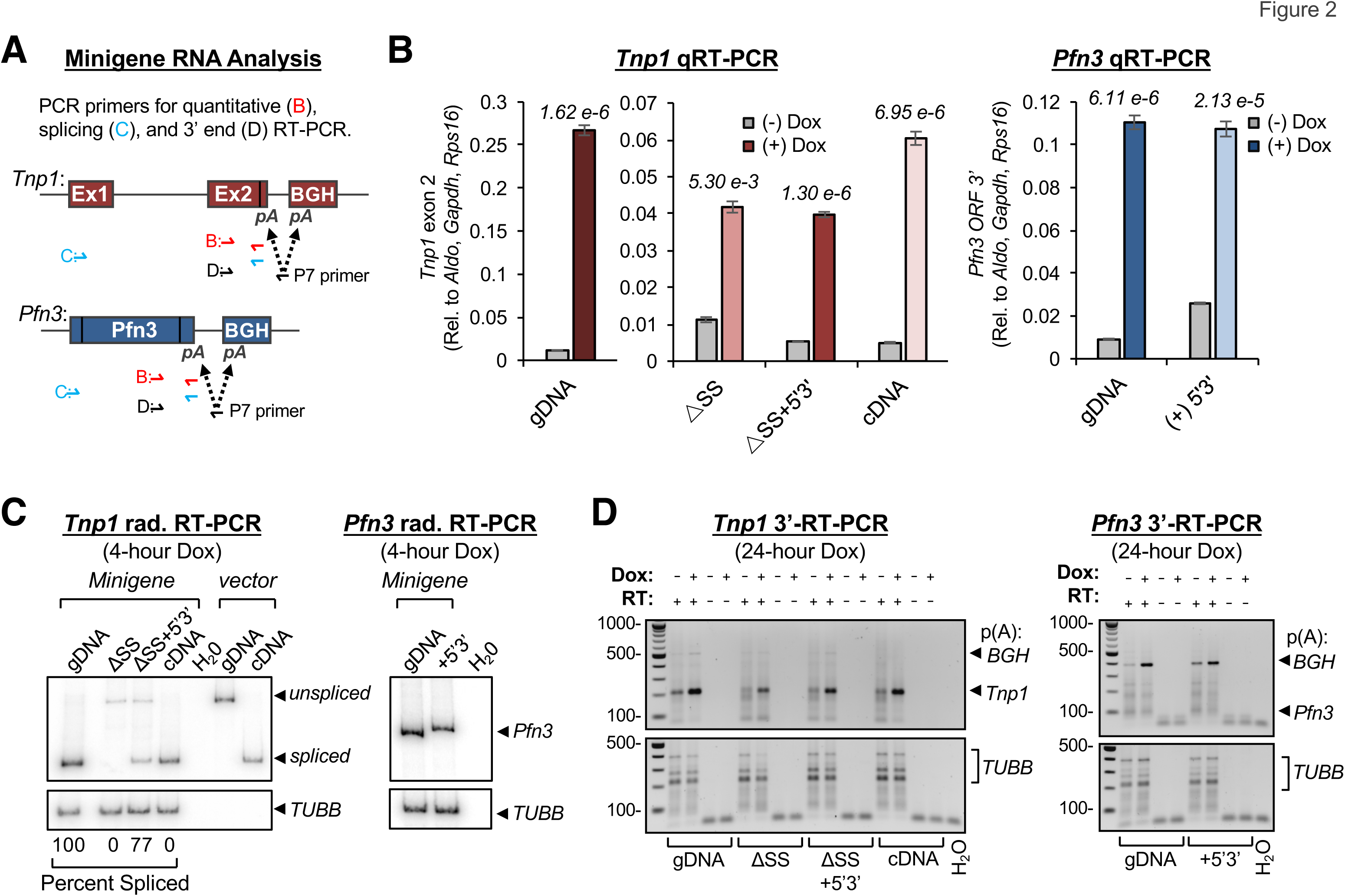
Analysis of minigene induction and splicing. **(A)** Schematic of primer target locations for analysis of minigene-derived RNA. **(B)** Quantitative RT-PCR (qRT-PCR) with primers directed against *Tnp1* exon 2 and the *Pfn3* ORF following 4 hours of treatment with Dox or vehicle. Minigene expression was normalized to the average of three reference genes (*ALDO, GAPDH*, and *RPS16*). Data shown are from three independent, biological replicates; error bars depict mean ± SEM. **(C)** Phosphorimaging analysis of 4 hour induced RT-PCR products generated with intron-flanking *Tnp1* primers or 5’ and 3’-directed *Pfn3* ORF primers in the presence of dCTP α^32^P. Percent-spliced-in (PSI) was determined as the signal intensity of the spliced product versus the sum of spliced and unspliced products. Amplification of *β-tubulin* (*TUBB*) was performed in a multiplex reaction as an internal control for loading (*bottom*). -**(D)** 3’-end RT-PCR to assess minigene polyadenylation. Distinct products reflect use of either the endogenous polyadenylation sites-encoded within the *Tnp1* and *Pfn3* minigenes or the vector-encoded site located within the BGH cassette. *β-tubulin* (*TUBB*) served as a loading control.

As biased chromatin signatures have been detected at the 3’ ends of genes (65), we further examined 3’-processing of the minigene-derived RNAs to ensure against secondary effects related to altered polyadenylation in response to splice site mutation. To achieve efficient 3’-processing, the minigenes were configured to retain their endogenous p(A) sites in addition to a downstream vector-encoded site (Fig. 1B). Precise p(A) utilization was assessed through RT with a hybrid P7 and anchored oligo(dT)_25_ primer, followed by PCR amplification with gene-specific forward primers and P7 reverse primer (Fig. 2A). Agarose gel electrophoresis confirmed that the minigene-derived transcripts undergo efficient 3’-processing and revealed a substantial increase in the polyadenylated fraction in response to Dox induction. Importantly, while 3’-RT-PCR uncovered distinctions in p(A) site preferences, splice site mutation did not alter site selection. For example, while the *Pfn3* minigenes preferentially utilized the vector-encoded site, *Tnp1* transcripts favored the endogenous p(A) site, irrespective of splicing status (Fig. 2D). Together, these results establish the isogenic minigene panel as a validated inducible resource for examining the influence of splicing on genic chromatin.

### Genome-wide depletion of H3K36me3 and 5mC at single- vs. multi-exonic genes

Having confirmed the expected expression patterns in the minigene panel, we next sought to determine whether genic DNA methylation is coupled to splicing-associated H3K36me3. However, to avoid confounding variables at the chromatin level, we first examined genome-wide data for additional modifications with evidence of splicing dependency. Drawing from the observation that H3K36me3 and DNA methylation are reduced at intronless genes (10-12), we surveyed for chromatin features that displayed differential accumulation at the gene bodies of expressed single-exon (unspliced) versus multi-exonic (spliced) genes. For this purpose, we selected human erythroleukemic K562 cells as a model system based on a wide availability of genome-wide data through the ENCODE project (66). Expressed genes were defined as single or multi-exonic based on gene annotations and were validated through examination of K562 RNA-seq data (Table S3). As select intragenic chromatin modifications show varied detection with increasing distance from the TSS, and single-exon genes are inherently shorter than their multi-exonic counterparts (67-69), genes were further segregated by length. Focusing on chromatin features associated with transcriptionally active loci (70), ChIP-seq analysis revealed generally decreased detection at single relative to multi-exonic genes. However, when genes of comparable size were considered, for the most part, the pattern of enrichment was not altered for unspliced versus spliced contexts: H3 lysine 4 trimethylation (K4me3), lysine 27 acetylation (K27Ac) and lysine 9 acetylation (K9Ac) peaked at the 5’ end of the gene followed by a steady decrease over the transcription unit, whereas lysine 79 dimethylation (K79me2) was elevated central to the gene body (Fig. 3A). When further considering transcription level, decreased detection of these modifications at single-exon genes showed a strong correlation to reduced pol II density: examination of promoter-proximal chromatin as a function of pol II levels showed the greatest enrichment of H3K4me3, H3K27Ac, H3K9Ac and H3K79me2 at the most highly transcribed genes, irrespective of splicing status (Fig. 3B).

**Figure 3.**
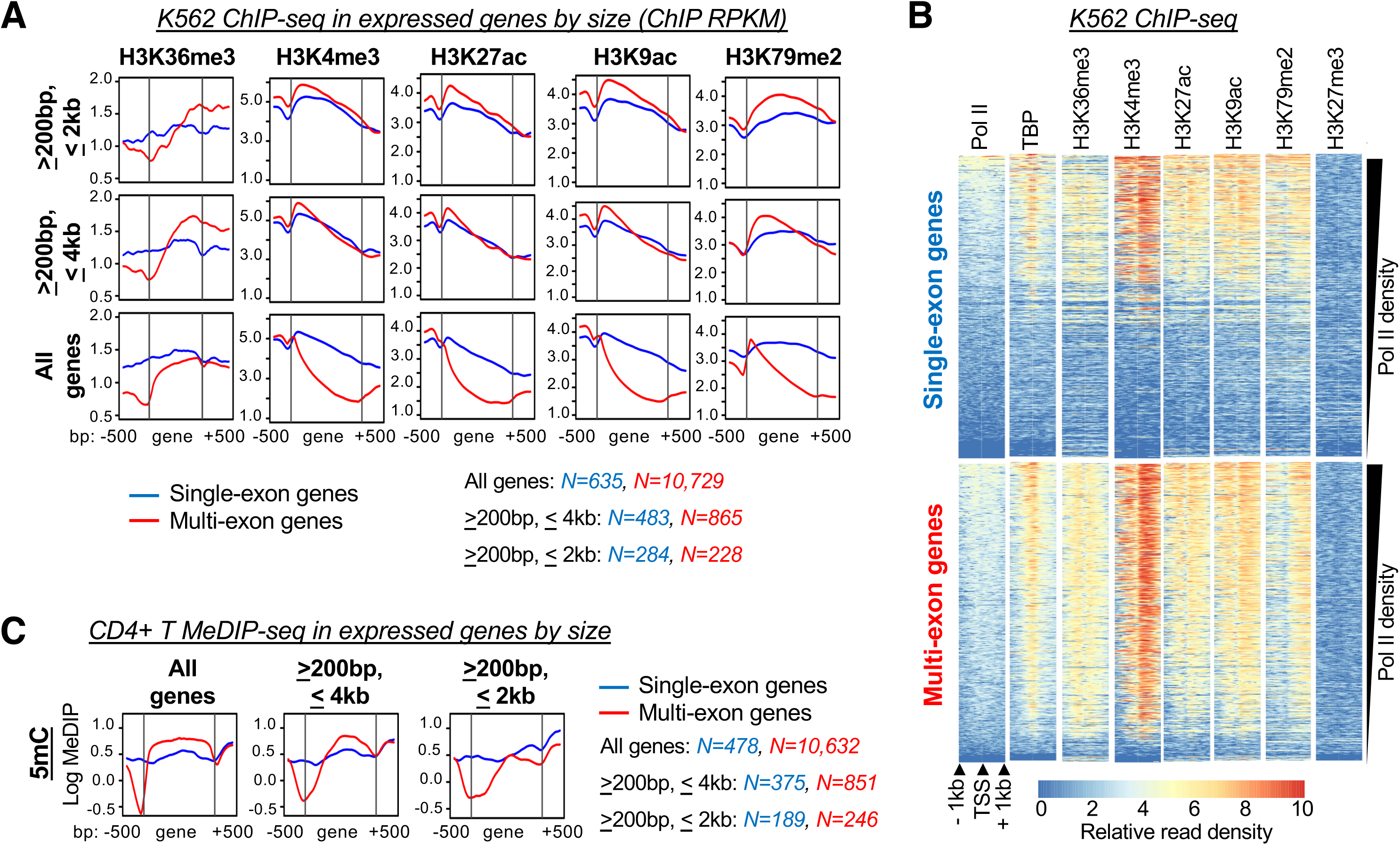
Epigenetic profiles associated with expressed single- vs. multi-exon genes in K562 and primary CD4+ human T cells. **(A)** Read coverage of transcription-associated histone post-translational modifications (PTMs) across the gene bodies of expressed genes in K562 cells. Genes were segregated into sets with different length restrictions; single- and multi-exon genes are shown in blue and red, respectively. **(B)** Heatmaps showing ChIP-seq read coverage for histone PTMs, pol II, and TATA binding protein (TBP) centered around transcription start sites (TSS) of single- and multi-exon genes; rows are sorted by decreasing pol II density. **(C)** Read coverage for 5-methylcytosine (5mC) MeDIP-seq in CD4^+^ T-cells over gene bodies of expressed genes. Read coverage is presented for all genes or segregated based on gene length. RPKM, reads per kilobase per million mapped (log scale); MeDIP, methylated DNA immunoprecipitation.

H3K36me3 represented a prominent exception amongst transcription-associated modifications, wherein detection over spliced genes gradually increased along the length of the coding unit while unspliced genes remained flat (Fig. 3A). This distinction was seen across all categories of gene length and was further observed in the pol II-stratified data, wherein highly transcribed single-exon genes often failed to accumulate H3K36me3 (Fig. 3B). These results are consistent with previous accounts of reduced H3K36me3 at intronless genes (10). To further examine whether genic 5mC follows a similar pattern of enrichment, we turned to our own genome-wide data performed in primary human CD4+ T cells (19). As in the K562 analysis, expressed single- versus multi-exonic genes were defined from gene annotations and RNA-seq analysis and segregated on the basis of length (Table S4). Examination of methylated DNA immunoprecipitation-sequencing (MedIP-seq) results established that 5mC was enriched across the gene body of spliced genes but, like H3K36me3, remained flat across unspliced genes of all sizes (Fig. 3C). While our current understanding of transcription-associated chromatin is not exhaustive, these findings reveal that the failure to accumulate H3K36me3 and DNA methylation at unspliced genes is relatively unique to these modifications.

### H3K36me3 is directly related to splicing in the 293 T-REx minigene panel

Based on the distinctiveness of H3K36me3 and 5mC in the genome-wide analysis of single versus multi-exonic genes, we returned to our minigene panel to explore the intra-dependency of these features as well as their relationship to splicing in an isogenic setting. As H3K36me3 was previously shown to exhibit splicing-dependency (10,25), we examined whether we could recapitulate this association within our minigenes. To this end, we pursued ChIP with a modification-specific antibody directed against H3 trimethylated at K36 and antibody to pan-H3 to normalize for variations in nucleosome density. Given the established reliance of H3K36me3 on pol II transcription (21), ChIP was performed in cells treated with vehicle or Dox for 24 hours to represent uninduced and induced states, respectively. Consistent with the observed low basal expression (Fig. 2B), the uninduced minigenes generally displayed minor gene body enrichment of H3K36me3 as compared to the surrounding sequence that for the most part showed no relationship to splicing capacity: H3K36me3 levels in the uninduced *Tnp1* gDNA, ΔSS, ΔSS+5’3’ and cDNA minigenes or *Pfn3* gDNA and +5’3’ minigenes were largely indistinct (Fig. 4A, 4B, dotted lines show basal gDNA H3K36me3). In contrast, examination of H3K36me3 in the induced minigenes showed a remarkable association with pre-mRNA splicing. H3K36me3 gains were directly related to splicing capacity in the two-exon isogenic panel, wherein the strongly spliced gDNA and poorly spliced ΔSS *Tnp1* minigenes showed the highest and lowest levels of H3K36me3, respectively, and the partially spliced ΔSS+5’3’ minigene fell between these two extremes (Fig. 4A). In fact, the induced splicing-deficient ΔSS and cDNA *Tnp1* minigenes exhibited a minor reduction in H3K36me3 that was not related to biased detection of overall H3 in response to gene activation (Fig. 4A, Fig. S3). A similar trend was observed in the single-exon minigenes, wherein both *Pfn3* constructs displayed a 3’ re-positioning of H3K36me3 in response to induction, but the introduction of heterologous splice sites in the *Pfn3*+*5’3’* minigene substantially increased the magnitude of H3K36me3 gains (Fig. 4B). Notably, while elevated H3K36me3 in the *Tnp1* gDNA minigene can be attributed to the higher level of induction, the remaining minigenes displayed comparable expression (Fig. 2B). Combined with the genome-wide analysis in single- vs. multi-exonic genes (Fig. 3A), these results establish that H3K36me3 is not strictly tied to transcription and solidifies a role for splicing in the modulation of this mark. Together, these data authenticate the isogenic minigene panel as an ideal substrate for examining the impact of splicing on genic DNA methylation through H3K36me3.

**Figure 4.**
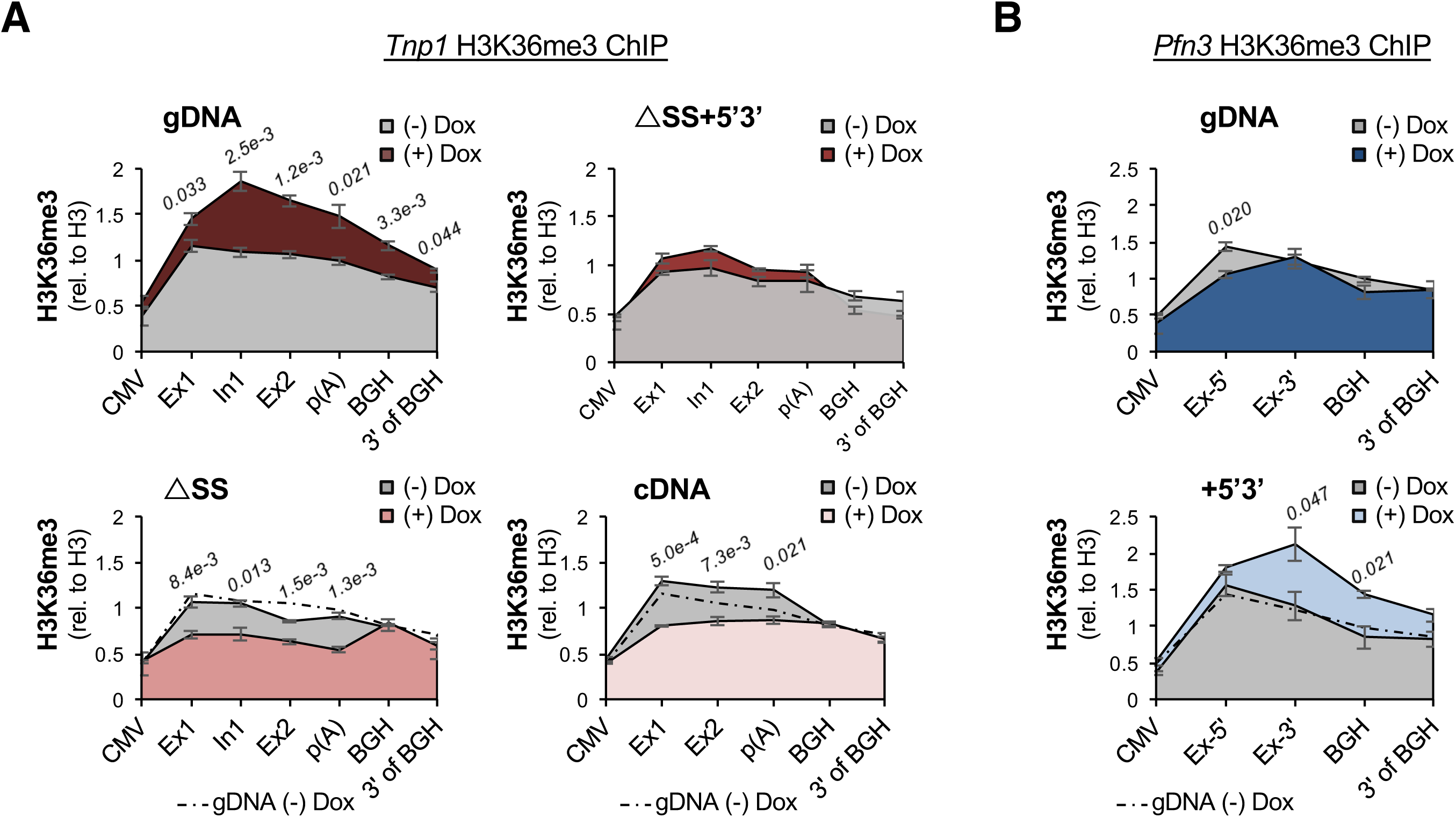
Induced H3K36me3 is associated with minigene splicing potential. **(A)** H3K36me3 ChIP and qPCR with primers against the *Tnp1* gene body and surrounding sequence in the *Tnp1* gDNA, ΔSS+5’3’, ΔSS, and cDNA minigenes relative to H3 ChIP at the analogous amplicons. H3K36me3 levels were assessed under basal conditions (*gray*) and following 24 hours of Dox induction (*red*). The dotted line overlay serves as a reference for the basal H3K36me3 in the gDNA minigene. **(B)** H3K36me3 ChIP-qPCR as in **(A)**, but with basal (*gray*) and induced (*blue*) *Pfn3* gDNA and +5’3’ minigenes. ChIP values represent mean ± SEM from three independent, biological replicates. *p-*values=two-tailed Student’s t-Test comparing the indicated amplicons (shown in italics).

### *De novo* DNA methylation is independent of splicing in the integrated minigenes

Armed with this well-characterized 293 T-REx inducible minigene panel representing distinct splicing capacities and H3K6me3 states, we turned to examining the splicing-dependency of genic DNA methylation. To assess the methylation status of the integrated *Tnp1* and *Pfn3* minigenes, we initially pursued methylation-specific restriction enzyme-PCR (MSRE-PCR). MSRE-PCR involves digestion of genomic DNA with isoschizomer restriction enzymes displaying distinct sensitivities to overlapping DNA methylation. Subsequent PCR amplification with primers that flank the restriction site allow for estimation of DNA methylation through increased resistance to cleavage in the methyl-sensitive as compared to insensitive digest (71,72). The sequences of the *Tnp1* and *Pfn3* minigenes and their derivatives naturally contain iterations of the tetranucleotide sequence 5’-CCGG-3’, which is recognized and cleaved by the isoschizomers HpaII and MspI (Fig. 5A). While MspI cleaves this sequence irrespective of methylation status, HpaII is blocked by methylation of the internal cytosine.

**Figure 5.**
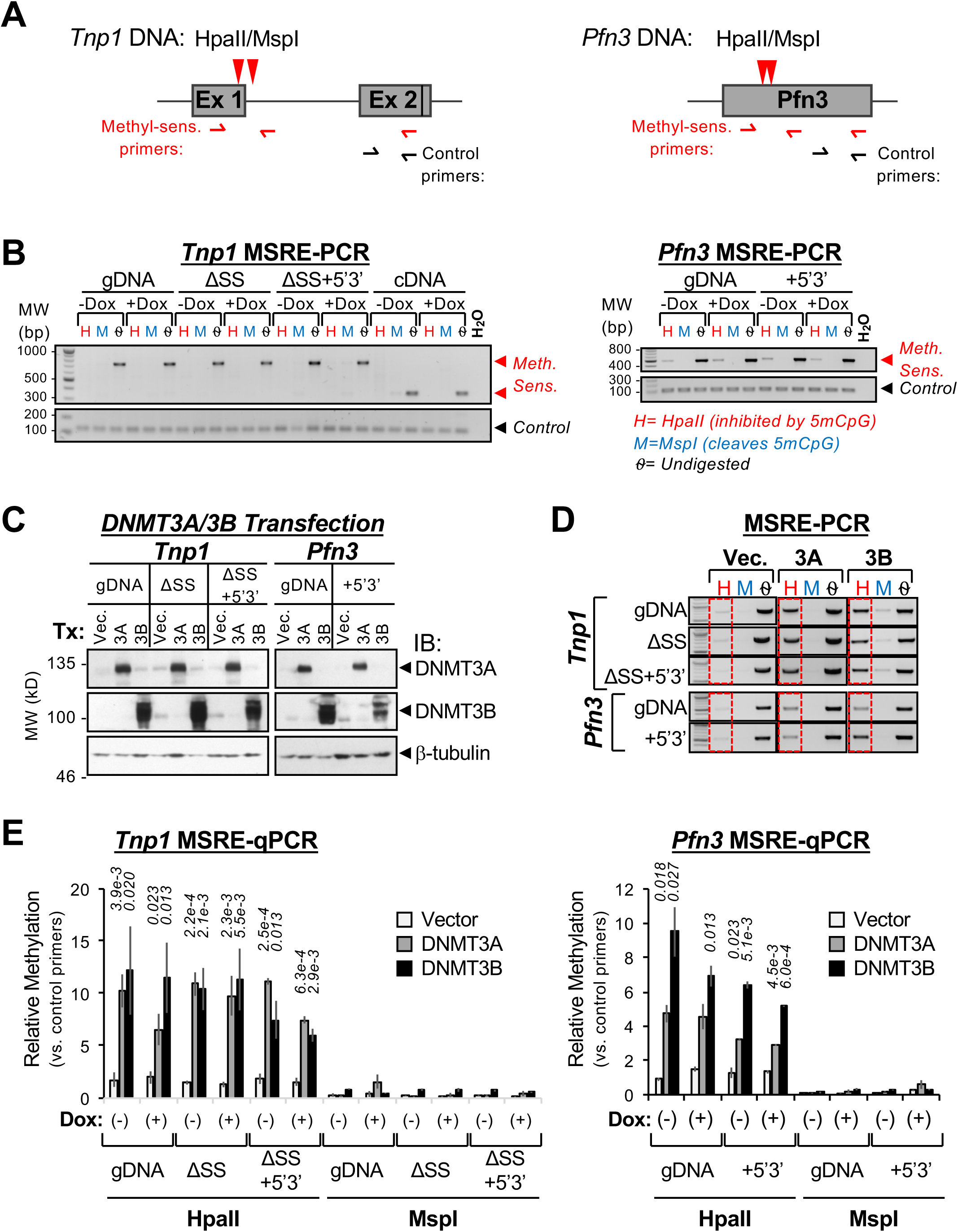
Detection of DNA methylation in *Tnp1* and *Pfn3* minigenes with DNMT3 overexpression. **(A)** Location of HpaII/MspI isochizomer target sequences in the *Tnp1* and *Pfn3* minigenes (red arrowheads). HpaII cleavage is inhibited by overlapping methylation, whereas MspI is insensitive. Primer binding sites flanking the restriction site (methyl-sensitive) and undigested control region are shown. **(B)** Methylation-sensitive restriction enzyme (MSRE)-PCR in basal and induced *Tnp1* and *Pfn3* minigenes. Minimal signal intensity in the HpaII-digested samples is indicative of minor to absent overlapping DNA methylation at the interrogated sites. **(C)** Immunoblotting for DNMT3 overexpression in *Tnp1* and *Pfn3* minigene host cells with transient transfection of mammalian expression constructs encoding DNMT3A or DNMT3B, as compared to vector control. Blotting for β-tubulin served as loading control. **(D)** MSRE-PCR in *Tnp1* and *Pfn3* host cells with overexpression of DNMT3A and DNMT3B. Gains in DNA methylation are evident through amplification with the methyl-sensitive primer sets following HpaII digest in DNMT3-transfected cells versus vector control (red boxes). (-) Dox samples shown; (+) Dox samples and control primers are presented in Figure S4. **(E)** MSRE-qPCR for quantification of resistance to HpaII digest (left), and concurrent susceptibility to MspI (right), in the DNMT3-transfected minigene host cells +/− Dox-induction. Values represent amplification with methyl-sensitive primers relative to the downstream control primer sets that do not overlap a HpaII/MspI site. Mean ± SD of two independent, biological replicates are shown. *p*-values=two-tailed Student’s t-Test comparing DNMT3A (left italic) and DNMT3B (right italics) to control. In addition, for *Pfn3*+5’3’ DNMT3A versus DNMT3B values were statistically significant (p=4.5e-3 and 5.0e-4 for minus and plus Dox, respectively).

In the case of *Tnp1*, the restriction sites directly interrogate methylation at the exon 1/intron 1 border, whereas the *Pfn3* sites are located centrally within the ORF (Fig. 5A). Initial assessment of DNA methylation in the minigene panel revealed basal hypomethylation that was not enhanced by transcriptional induction. MSRE-PCR across the entire set of uninduced and induced *Tnp1* minigenes showed little, if any, protection against HpaII (methyl-sensitive) as compared to MspI (methyl-insensitive) cleavage (Fig. 5B, *top*). PCR amplification with primers directed to exon 2, which lacks HpaII/MspI recognition sequences, served as loading control and demonstrated the integrity of the genomic DNA samples following extended digestion (Fig. 5B, *bottom*). Similarly, while some HpaII resistance was observed at the *Pfn3* minigenes, the overall level was rather low and was not increased by induction of transcription (Fig. 5B, *right*). These findings indicate that elevated H3K36me3 over the gene bodies of the expressed and spliced minigenes was not sufficient to promote genic DNA methylation in the context of 293 T-REx cells.

Notably, expression of the PWWP-containing *de novo* DNA methyltransferases is generally low in peripheral cells (73), raising the possibility that the determined minigene hypomethylation relates to inadequate DNMT3 expression. Indeed, immunoblot revealed low expression of DNMT3A and DNMT3B across the isogenic 293 T-REx panel (Fig. 5C). This observation fortuitously allowed us to layer DNA methylation into our minigene cell lines through exogenous introduction of the *de novo* DNA methyltransferases (Fig. 5C). To this end, we performed transient transfection of DNMT3A along with its regulatory factor DNMT3L, or myc-tagged DNMT3B in *Tnp1* and *Pfn3* minigene host cells prior to Dox induction. DNMT3 overexpression was confirmed through immunoblot as compared to vector control (Fig. 5C, Fig. S4A). To avoid complications related to overall minigene length and GC content, both of which could impact DNA methylation through H3K36me3, the intron-deleted *Tnp1* cDNA minigene was omitted from further analyses. As a reminder, the *Tnp1* ΔSS minigenes employed conservative nucleotide substitutions that preserved CG density and the *Pfn3*+*5’3’* minigene was solely modified through the introduction of short heterologous splice sites flanking the ORF (Fig. 1B). Strikingly, MSRE-PCR revealed a robust increase in resistance to digestion by methylation-sensitive HpaII in the DNMT3 transfected cells relative to control (Fig. 5D, *red boxes*). This increase was observed in response to both DNMT3A and DNMT3B and was seemingly independent of minigene expression: similar gains in DNA methylation were detected in the presence or absence of subsequent Dox-induction (Fig. 5D, Fig. S4B-C). Importantly, elevated DNA methylation was seen across the entire panel of *Tnp1* and *Pfn3* minigenes, including the splicing-deficient *Tnp1* ΔSS and *Pfn3* clones, which failed to accumulate H3K36me3 in response to induction (Figs. 2B, 4A, 4B). Quantitative-PCR analysis of the MSRE-digested DNA further confirmed significant gains in DNA methylation across the minigene panel, as assessed through amplification with restriction site-flanking primers relative to control in the HpaII and MspI treated samples (Fig. 5E). Curiously, DNMT3A was less effective at methylating the interrogated sites in the *Tnp1* gDNA and *Pfn3* minigenes. Nevertheless, HpaII/MspI digest clearly established efficient methylation of the *Tnp1* and *Pfn3* minigenes, including in the absence of transcriptional induction and pre-mRNA splicing (Fig. 5E).

A main drawback of the MSRE-qPCR approach is the limited availability of methyl-sensitive restriction sites. As we were primarily interested in examining the impact of H3K36me3 on DNA methylation, and trimethylation of H3K36 is enriched towards the 3’ end of genes, we additionally pursued high-resolution analysis through BS-pyrosequencing in the Dox-induced minigenes (Fig. 6A). Like traditional bisulfite-sequencing, BS-pyrosequencing relies on chemical conversion of unmethylated cytosines to uracil, while leaving methylated cytosines intact. DNA sequencing then allows for precise determination of percent methylation at interrogated cytosines through quantification of pyrophosphate release during stepwise deoxyribonucleotide triphosphate (dNTP) incorporation (74). Assays were designed to examine the 7 CpGs that overlap *Tnp1* exon 2 and 18 CpGs found within the *Pfn3* ORF (Fig. 6A). Consistent with the MSRE-qPCR results, BS-pyrosequencing in the DNMT3A- and 3B-transfected *Tnp1* and *Pfn3* minigene host cells showed efficient *de novo* methylation across the queried CpGs (Figs. 6B, 6C). While levels at specific CpGs showed some variability, average gains in methylation of 10-15% were observed in the DNMT3- transfected cells that were generally unrelated to splicing capacity (Figs. 6B, 6C, *left* and *right*, respectively). *Tnp1* gDNA transfected with DNMT3A represented an exception, wherein methylation was reduced relative to the splicing-deficient minigenes (Fig. 6B). This result is consistent with the MSRE-qPCR result directed against the 5’ end of the coding unit, suggestive of a *bona fide* defect (Fig. 5E). In contrast, the site-specific reduction in DNMT3A-directed methylation of the *Pfn3* minigenes determined through MSRE-qPCR was not observed in the more comprehensive pyrosequencing results (Figs. 5E, 6C). Overall, analysis of DNA methylation in the DNMT3- transfected 293 T-REx cells revealed efficient *de novo* methylation of the integrated minigene DNA, irrespective of splicing status. As the set of minigenes characterized by competent splice sites (*Tnp1* gDNA/ΔSS +5’3’ and *Pfn3*+5’3’) showed elevated H3K36me3 post-induction as compared to their counterparts lacking functional splice sites (*Tnp1* ΔSS and *Pfn3* gDNA) (Figs. 4A, 4B), these findings further indicate that trimethylation of H3K36 is not an obligate platform for the recruitment of DNMT3 enzymes at transcribed genes. Based on the sum of these minigene data, we conclude that variations in DNA methylation that occur between spliced and unspliced genes are not solely related to splicing-associated H3K36me3.

**Figure 6.**
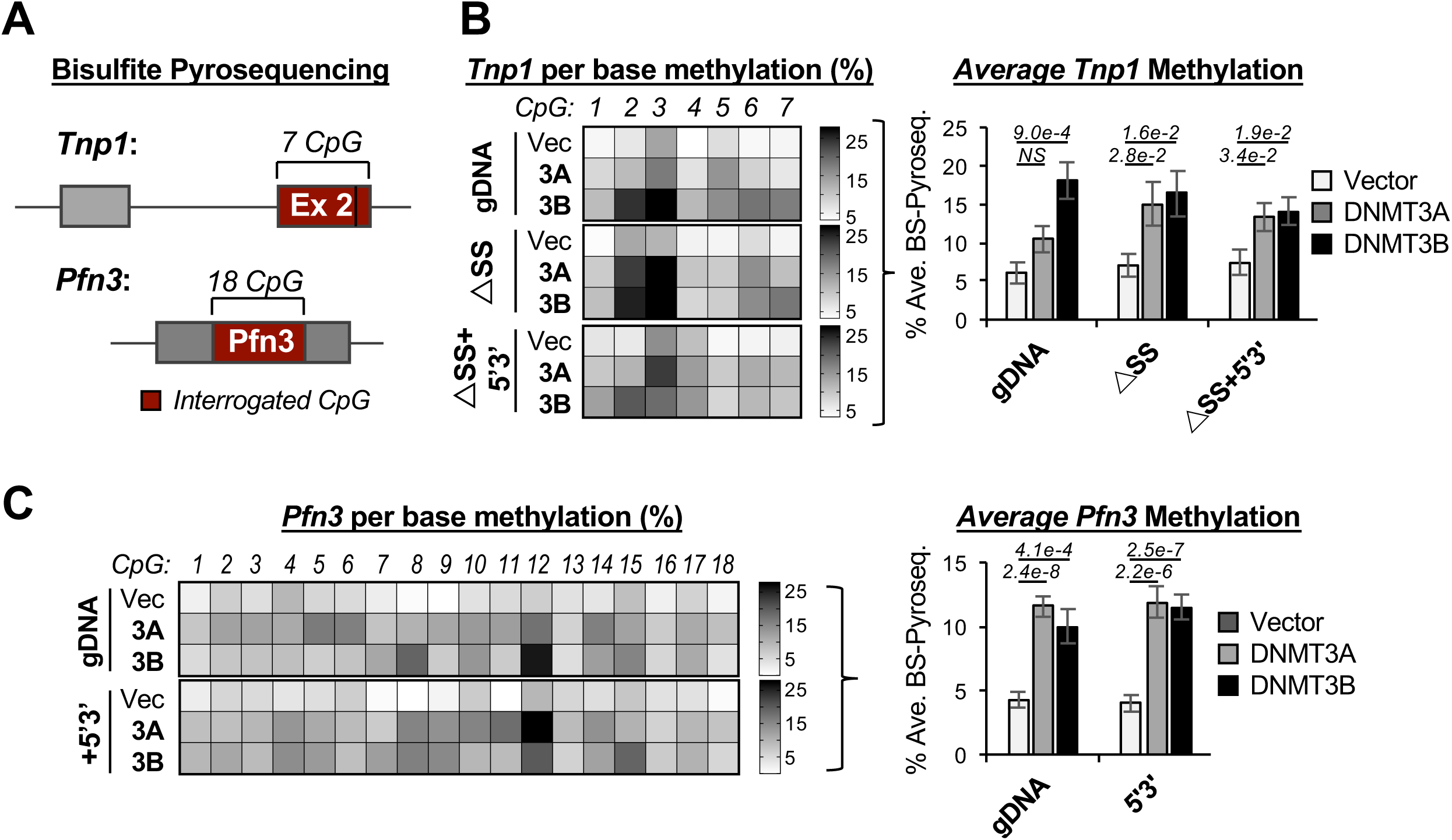
Base-resolution determination of CpG methylation in *Tnp1* and *Pfn3* minigenes with exogenous DNMT3 expression. **(A)** Schematic representation of regions interrogated by bisulfite-pyrosequencing (BS-pyroseq) in the *Tnp1* and *Pfn3* minigenes. **(B)** Per-base heatmap for percent CpG methylation at the 7 interrogated sites within *Tnp1* exon 2. Values represent averages from duplicate BS-pyroseq performed on biological replicates of Dox-induced *Tnp1* host cells transfected with vector encoding DNMT3A, DNMT3B or control (left). Average methylation across the 7 sites in DNMT3-transfected and control cells (right), mean ± SEM. **(C)** BS-pyroseq of the 18 CpGs overlapping the *Pfn3* ORF, as in **(A)**. *p*-values=two-tailed Student’s t-Test comparing average methylation DNMT3-transfected cells versus control, as indicated.

## DISCUSSION

Recent investigations support a role for local modulation of intragenic chromatin structure in the regulation of pre-mRNA splicing (13,75-78). However, little is known of the mechanisms supporting asymmetric distribution of chromatin to gene bodies. Variations in chromatin structure exist between distinct genes and within single genes, wherein select modifications are differentially detected at exons versus introns. In particular, H3K36me3 and DNA methylation are enriched at expressed exons and are reciprocally depleted at intronless genes (10-12,65,79) (Fig. 3). These observations at the genomic level call into question the mechanism by which the responsible methyltransferases discriminate between spliced and unspliced genes. As a central evaluator of splicing potential, here we explore the role of the spliceosome itself in the establishment of genic chromatin. To this end, we generated a panel of minigenes representing distinct splicing states for stable integration into a single FRT site in 293 T-REx cells. Minigene expression is driven by an inducible promoter in this system, thus uncoupling the respective impacts of transcription and pre-mRNA splicing on genic chromatin. With the specific goal of examining the splicing-dependency of DNA methylation and potential relationship to H3K36me3, we observe an unexpected degree of independence between these parameters.

As a universal hallmark of transcription, H3K36me3 has been heavily interrogated in a variety of model systems. The diverse functions attributed to H3K36me3 include the prevention of cryptic transcription, regulation of pre-mRNA splicing through recruitment of RNA binding proteins, and an emerging role in the DNA damage response (13,16,26,80-86). Appropriately, the deposition of H3K36me3 within gene bodies is equivalently nuanced and is subject to regulation through both the transcriptional and splicing machineries. The responsible methyltransferase, SETD2, is targeted to expressed genes through direct interaction with elongating pol II and this ability is compromised upon inhibition of pre-mRNA splicing: chemical interference of splicing and splice site mutations result in reduced SETD2 and H3K36me3 at affected gene bodies (10,23,25) (Fig. 4). However, the mechanistic basis of H3K36me3 splicing-dependency remains unknown. Despite an extensive body of biochemical literature, to our knowledge, direct interaction between SETD2 and spliceosomal components has not been reported. In addition, SETD2 homologs are efficiently targeted to transcribed genes in organisms that engage in little to no splicing, thus supporting a splicing-independent component (26). These somewhat contradictory observations underscore the inherent complexity of intragenic chromatin modifications in organisms that shape their chromatin landscapes in response to transcription and splicing. Remarkably, we recapitulated this dual dependency within our minigene panel, wherein trimethylation of H3K36 was responsive to both transcription and splicing (Figs. 2, 4). Furthermore, the level of H3K36me3 gains directly reflected splicing potential, such that mutations that partially or fully ablated splicing of the two-exon minigene resulted in comparable decreases in induced H3K36me3 (Figs. 2C, 4A). Reciprocally, addition of strong splice sites to the single-exon minigene produced a moderate increase in H3K36me3 as compared to the unspliced parental minigene (Fig. 4B). Given that the orientation of the incorporated sites generated a poor substrate for the spliceosome, the associated increase in H3K36me3 in the *Pfn3*+*5’3’* minigene suggests that the consensus sequences are indeed recognized by SETD2 or associated factors at the chromatin template through a currently unknown mechanism. The notion that splicing factors function in the regulation of transcription or chromatin, independently of pre-mRNA splicing, is supported through accounts of splicing proteins moonlighting in transcriptional complexes and through the prevalence of spliceosomal proteins in organisms with few introns (87-91). In sum, the generated minigene panel recapitulates the observed genome-wide increase in H3K36me3 at multi-versus single-exon genes and further establishes that transcription is not sufficient to promote genic H3K36me3 in the absence of splicing signals (Fig. 4, Fig. S3, Fig. 7).

**Figure 7.**
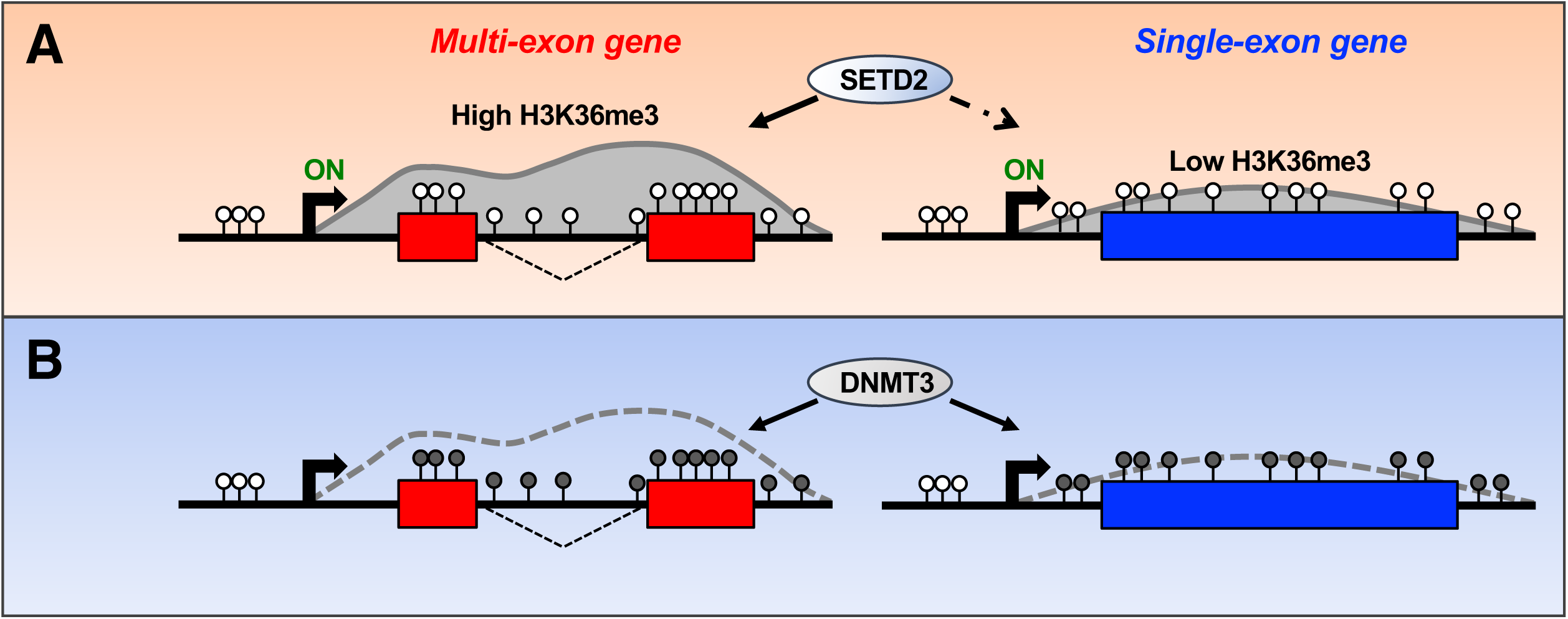
Summary model of current findings. **(A)** Pre-mRNA splicing signals promote SETD2-catalyzed H3K36me3 at actively transcribed gene-bodies. **(B)** The DNMT3 enzymes efficiently methylate cytosines in CpG dinucleotides overlapping gene bodies irrespective of transcriptional status, H3K36me3 levels or pre-mRNA splicing capacity.

While the observations related to H3K36me3 in our minigene panel were intriguing in their own right, our main goal was to determine the relevance of splicing and H3K36me3 to the generation of intragenic DNA methylation. Given that the distribution of DNA methylation within gene bodies closely mirrors H3K36me3 and the *de novo* DNA methyltransferases contain a H3K36me3-interacting PWWP domain, we asked whether splicing-dependent trimethylation of H3K36 functions as a platform for DNMT recruitment to spliced genes (10-12,25,36,37). In support of such a platform, ablation of either *SETD2* or disruption of the DNMT3B PWWP domain in mouse embryonic stem cells (mESCs) led to decreased gene body DNA methylation at transcribed genes (39). Separately, exogenous introduction of DNMT3B into the budding yeast *S. cerevisiae* resulted in H3K36me3-dependent increases in DNA methylation (38). However, considering that H3K36me3 is omnipresent at transcribed genes in eukaryotes, including at unspliced genes in *S. cerevisiae*, the determined changes in cytosine methylation upon DNMT3 modulation were relatively modest (26,38,39). Combined with accounts that cytosine methylation is concentrated at select exons in organisms with overall low DNA methylation, these findings argue against a simplistic H3K36me3-directed recruitment model (15,32,33). Accordingly, we observed substantial independence between H3K36me3 and cytosine methylation in our minigene cell lines. Exogenous introduction of DNMT3A or DNMT3B into basally hypomethylated 293 T-REx host cells led to comparable gains in DNA methylation in the splicing-deficient minigenes with low H3K36me3 and the splicing-competent minigenes with elevated H3K36me3, in the presence or absence of induction (Figs. 4-6). While some variability was observed at specific CpGs, overall increases in methylation were roughly comparable, reaching a maximum of 20-25% at select sites in both the spliced and unspliced constructs (Fig. 6B, 6C). Curiously, introduction of DNMT3A into 293 T-REx cells harboring the *Tnp1* gDNA minigene represented a sole exception wherein CpG methylation across the minigene was attenuated as compared to the analogous splicing-deficient constructs (Figs. 5E, 6B). The reason for this reproducible decrease is unclear, but it is worth noting that the DNMT3A enzyme was less adept at methylating genic CpGs when re-introduced into knockout mESCs as compared to DNMT3B (39). Considering that the strongly spliced *Tnp1* gDNA minigene intuitively represented the most likely substrate for DNMT3 activity based on the genomewide elevation of DNA methylation at spliced genes (11,14,15) (Fig. 3C), the combination of our current results and published observations may reflect an unexpected inhibitory role for splicing in the regulation of DNMT3A function.

While our minigene results in 293 T-REx cells establish that ablation of pre-mRNA splicing impacts H3K36me3 without compromising *de novo* DNA methylation, they do not provide a framework as to why DNA methylation positively correlates with pre-mRNA splicing in genome-wide data. A trivial explanation would relate to reduced CpG density at unspliced locations. However, elevated DNA methylation at exons versus introns persists after correcting for CpG density (35), arguing against a simplistic sequence bias. We envision several scenarios that may distinguish our integrated unspliced minigenes from their endogenous counterparts. For one, the entire panel of minigenes is incorporated into a single genomic location. It is possible that unspliced genes are naturally arrayed at loci that are ineffective substrates for the DNMT3 enzymes in their normal chromosomal context. Alternatively, perhaps unspliced genes efficiently acquire *de novo* methylation, but are less able to maintain it through reduced DNMT1 activity post-replication or increased TET-catalyzed demethylation (28,92). Unfortunately, the 293 T-REx system utilized in this study does not allow for examining the relationship between H3K36me3 and DNA methylation in a maintenance context, as minigene expression is rapidly silenced following Dox removal (93). While ideal for uncoupling the respective impacts of transcription and splicing on the assessed chromatin features in the current work, sustained minigene expression will ultimately be required to determine whether splicing guides genic DNA methylation in a constitutively expressed context.

In sum, we describe the generation of a minigene resource that unequivocally demonstrates independence between pre-mRNA splicing and *de novo* DNA methylation in an isogenic chromosomal context, wherein splicing-deficient and –competent substrates were equivalently targeted by the DNMT3 enzymes. In contrast, and consistent with previous reports, splicing positively correlated with H3K36me3 and transcriptional induction within the panel (10,25,60,94-96) (Fig. 2, 4). This latter demonstration is particularly important in that it establishes that the minigenes accurately recapitulate features of endogenous genes in an adaptable heterologous context. While the current work specifically addresses the impact of splicing on DNA methylation, the panel was designed to function as a flexible platform to examine coupling between several molecular machineries.

Considering the numerous described connections between pre-mRNA splicing, transcription, genic chromatin and mRNA export (10,13,18,19,25,67,97-101), a better understanding of how these processes coordinate to regulate gene expression is clearly warranted. This is of particular relevance given that a number of critical human genes are of the single-exon variety, including the genes encoding c-Jun and the type I interferons (102,103). Deeper investigations within the minigene panel may reveal the principles by which single-exon genes bypass regulatory elements that otherwise act to reduce expression when splicing signals are removed from multi-exonic cassettes. Beyond the influence of splicing on transcription and genic chromatin, how distinct chromatin states impact pre-mRNA processing can be examined through CRISPR/Cas9 technology aimed at targeting chromatin-modifying enzymes to distinct sequences (104). On a related note, the presence of dual competing polyadenylation signals allows for investigation into the mechanisms governing alternative polyadenylation. All in all, the isogenic 293 T-REx minigene panel represents a diverse platform for dissecting associations between molecular processes, as we evidence here in finding that *de novo* DNA methylation displays independence from downstream pre-mRNA splicing.

## ACKNOWLEDGEMENT

We thank the members of the Advanced Technology Research Facility (ATRF) at the National Cancer Institute (Frederick) for providing bisulfite-pyrosequencing services. This study utilized the high-performance computational capabilities of the NIH HPC Biowulf cluster at the National Institutes of Health, Bethesda, MD (http://hpc.nih.gov).

